# CAR T-cell therapy - Paving the way for sensitized kidney transplant patients

**DOI:** 10.1101/2023.08.24.554644

**Authors:** Tambi Jarmi, Yan Luo, Rose Mary Attieh, Yaqing Qie, Martha E. Gadd, Tanya Hundal, Shennen Mao, Hemant S. Murthy, Burcin C. Taner, Mohamed A. Kharfan-Dabaja, Hong Qin

**Author notes:** **Correspondences:** Hong Qin, MD, PhD, Associate Professor of Medicine, Division of Hematology/Oncology, Director, Regenerative Immunotherapy & CAR-T Translational Research Program, Mayo Clinic Florida, 4500 San Pablo Rd S, Jacksonville, FL 32224, Tambi Jarmi, MD (co-correspondence), Assistant Professor of Medicine, Division Chair, Transplantation Medicine, Department of Transplantation, Mayo Clinic Florida, 4500 San Pablo Rd S, Jacksonville, FL 32224. These authors have made equal contributions to the project (Tambi Jarmi, Yan Luo).

## Abstract

Anti-HLA donor specific antibodies have been extensively documented for their critical role in kidney transplant rejection and resulting adverse outcomes. Several approaches have been employed to desensitize these patients; however, none of these explored therapeutic approaches has exhibited enduring clinical benefits. In this study, we explore a novel strategy of utilizing chimeric antigen receptor T cells (CAR T-cells) to target B cells in sensitized kidney transplant recipients. Specifically, we investigate the potential of our innovative MC10029 CAR T-cells, which are designed to recognize the B cell activating factor receptor (BAFF-R). BAFF-R is predominantly expressed on mature B cells and plays a crucial role in their survival, as well as in the promotion of autoreactive B cell. Our data revealed that sensitized patients’ B cells exhibited high levels of BAFF-R expression. We have successfully generated patient-derived MC10029 CAR T-cells from 6 sensitized patients. All these patient-derived MC10029 CAR T-cells consistently exhibited antigen-specific cytotoxicity against autologous B cells, accompanied by the release of cytotoxic granules. We have recently obtained FDA approval of an Investigational New Drug application for MC10029 CAR T-cell therapy in B-cell hematological diseases. This significant milestone paves the way for the pioneering launch of a human clinical trial, marking the first-ever application of CAR T-cell therapy in sensitized patients waiting for life-saving organ transplants.

## INTRODUCTION

Organ transplant demands, especially among patients with End Stage Kidney Disease (ESKD), far surpass the current availability of organs. This challenging situation is further compounded by the presence of pre-transplant high levels of anti-HLA antibodies, clinically known as sensitization^1^, making the matching process a highly complex one. Consequently, the waiting time on a transplant waitlist can be substantially lengthy, resulting in dire consequences where patients may tragically succumb to organ failure before receiving a life-saving organ transplant.^2^ To address this issue, the Organ Procurement and Transplant Network (OPTN) implemented a revised Kidney Allocation System (KAS) in 2014 to achieve a greater balance of utility and equity in deceased donor kidney allocation and to improve the rates of deceased donor kidney transplants (DDKT), specifically for sensitized patients with high calculated panel reactive antibody (cPRA). Unfortunately, despite KAS, the most highly sensitized candidates with >99.5% PRA still do not get their fair share of transplants.^3, 4^ Several approaches have been employed to desensitize these patients, aiming at reducing the burden of alloantibodies. These include plasmapheresis, the use of monoclonal antibodies that target B cells and the complement system, as well as the application of IgG-degrading enzyme to cleave HLA donor-specific antibodies.^5^ Despite these efforts, none of these therapeutic approaches have exhibited enduring clinical benefits.^6^ In this study, we explore a novel strategy of utilizing chimeric antigen receptor T cells (CAR T-cells) to target and eliminate B cells in sensitized kidney transplant recipients. Specifically, we investigate the potential of our innovative MC10029 CAR T-cells, which are designed to recognize the B cell activating factor receptor (BAFF-R). BAFF-R is predominantly expressed on mature B cells and plays a crucial role in their survival, as well as in the promotion of autoreactive B cell.^7^

## RESULTS

### Study Design

The primary objective of this study is to investigate the cytotoxicity of patient-derived MC10029 CAR T-cells against their own autologous B cells. A total of 10 eligible patients were recruited for this research, following the guidelines outlined in an approved Institutional Review Board protocol (IRB# 22-007826). Inclusion criteria consisted of adult patients aged 18 and above, exhibiting a high sensitization level with a calculated panel reactive antibody (cPRA) of 98% or higher, and expressing their willingness to provide informed consent, clinical data, and blood samples. Patients who had received desensitization therapy were excluded from the study.

Patient- and disease-related characteristics of these subjects are presented in **Table 1**. A unique and optimized protocol was developed, enabling the generation of patient-derived CAR T-cells from minimal amount of blood samples (**Supplementary Figure S1**). Additionally, the enriched B cells from the same blood samples were utilized as target cells in the presented CAR T-cell functional experiments (**Supplementary Methods**). Six out of the 10 patient samples provided an adequate quantity of both T cells and B cells necessary for performing the designed autologous experiments.

**Table 1:** Patient characteristics and responses to incubation of autologous B cells with either autologous non-CAR T or MC10029 CAR T-cells as determined by CAR T-cell cytotoxicity markers of CD107a and granule protein release^a^.

|  | **Patient 1** |  | **Patient 2<sup>b</sup>** |  | **Patient 3** |  | **Patient 4** |  | **Patient 5** |  | **Patient 6** |  |
| --- | --- | --- | --- | --- | --- | --- | --- | --- | --- | --- | --- | --- |
| **Age (years)** | 63 |  | 70 |  | 52 |  | 25 |  | 51 |  | 34 |  |
| **Gender** | Male |  | Female |  | Female |  | Female |  | Female |  | Male |  |
| **Race** | African American |  | Caucasian |  | African American |  | Caucasian |  | African American |  | Caucasian |  |
| **Disease** | Diabetic Nephropathy/ allograft failure |  | P-ANCA vasculitis |  | FSGS/allograft failure |  | FSGS/allograft failure |  | FSGS/allograft failure |  | Microscopic polyangiitis/ allograft failure |  |
| **Previous Organ Transplant** | Yes |  | No |  | Yes |  | Yes |  | Yes |  | Yes (twice) |  |
|  | **Non CAR-T** | **CAR-T<sup>c</sup>** | **Non CAR-T** | **CAR-T<sup>c</sup>** | **Non CAR-T** | **CAR-T<sup>c</sup>** | **Non CAR-T** | **CAR-T<sup>c</sup>** | **Non CAR-T** | **CAR-T<sup>c</sup>** | **Non CAR-T** | **CAR-T<sup>c</sup>** |
| **Degranulation (CD107a %)<sup>d</sup>** | 3.4 | **33.3** | ND | **51** | 3.83 | **31.1** | 4.53 | **13.1** | ND | **14.9** | ND | **49.8** |
| **Granzyme A** |  |  |  |  |  |  |  |  |  |  |  |  |
| **pg/ml** | 583 | **3310\*\*** | 746 | **8052\*\*** | 403 | **2500\*\*** | 270 | **2313\*\*** | 3373 | **9252\*\*** | 299 | **2564\*\*** |
| **Fold change<sup>e</sup>** | 1 | **5.7** | 1 | **10.8** | 1 | **6.2** | 1 | **8.6** | 1 | **2.7** | 1 | **8.6** |
| **Granzyme B** |  |  |  |  |  |  |  |  |  |  |  |  |
| **pg/ml** | 385 | **1188\*\*** | 154 | **2587\*\*** | 89 | **447\*\*** | 606 | **1625\*\*** | 289 | **889\*\*** | 456 | **2655\*\*** |
| **Fold change<sup>e</sup>** | 1 | **3.1** | 1 | **16.8** | 1 | **5.0** | 1 | **2.7** | 1 | **3.1** | 1 | **5.8** |
| **Perforin** |  |  |  |  |  |  |  |  |  |  |  |  |
| **pg/ml** | 1064 | **3690\*\*** | 716 | **3756\*\*** | 524 | **2234\*\*** | 399 | **1857\*\*** | 1788 | **4650\*** | 790 | **3063\*\*** |
| **Fold change<sup>e</sup>** | 1 | **3.5** | 1 | **5.3** | 1 | **4.3** | 1 | **4.7** | 1 | **2.6** | 1 | **3.9** |

ND = Not determined; Insufficient PBMCs required preferred production of MC10029 CAR T-cells and did not allow for both degranulation and secreted protein determination via ELISA; P-ANCA, Perinuclear anti-neutrophil cytoplasmic antibodies; FSGS, Focal segmental glomerulosclerosis
<sup>a</sup> The levels of granule proteins are presented as means from a total of 3 or 4 replicates, the P values were measured for the comparison between CAR-T and Non CAR-T (\* p< 0.05; \*\* p< 0.01).
<sup>b</sup> Patient 2 received prior treatment with rituximab in 2020 and cyclophosphamide until 2022.
<sup>c</sup> CAR-T = MC10029 CAR T-cells
<sup>d</sup> CD107a surface expression % is determined by flow cytometry
<sup>e</sup> Fold change is calculated by dividing pg/ml from MC10029 CAR T-cells by those from non-CAR T-cells.

### Targeting BAFF-R on patients’ B cells by MC10029 CAR T-cells

The confirmation of enriched B cells from patient blood samples was validated through flow cytometry analysis, using healthy donor PBMCs as controls for both T and B cell immunostaining. This enrichment process efficiently depleted CD3-positive T cells while enriching CD20-positive B cells. Notably, sensitized patients’ B cells exhibited high levels of BAFF-R expression (**Figure 1a**), highlighting the potential of targeting BAFF-R for B cell depletion in desensitization strategies. The cytotoxicity of patient-derived MC10029 CAR T-cells against autologous B cells was assessed by evaluating their degranulation activity and further confirmed through the measurement of granzyme B release. **Figure 1** illustrates representative patient data, demonstrating the readily detectable degranulation activity of MC10029 CAR T-cells in response to autologous B cells (**Figure 1b**). Furthermore, the CAR T-cells exhibited a significant release of cytotoxic granzyme B upon encountering autologous B cells, in comparison to non-CAR T-cells derived from the same patient. The non-CAR T-cells served as a control in both degranulation assay and granzyme B release experiment, representing the baseline level of T cell activation observed in each individual patient (**Figure 1c**). Importantly, the CAR T-cell functionality was shown to be antigen-specific, as evidenced by the activation of CAR T-cells exclusively against Nalm-6 cells expressing endogenous BAFF-R, while showing no activation against their genetically engineered variants lacking BAFF-R expression (**Figure 1b-c**). These data validate our developed autologous experimental system and support our hypothesis of utilizing MC10029 CAR T-cell therapy as a novel desensitization strategy.

**Figure 1.**
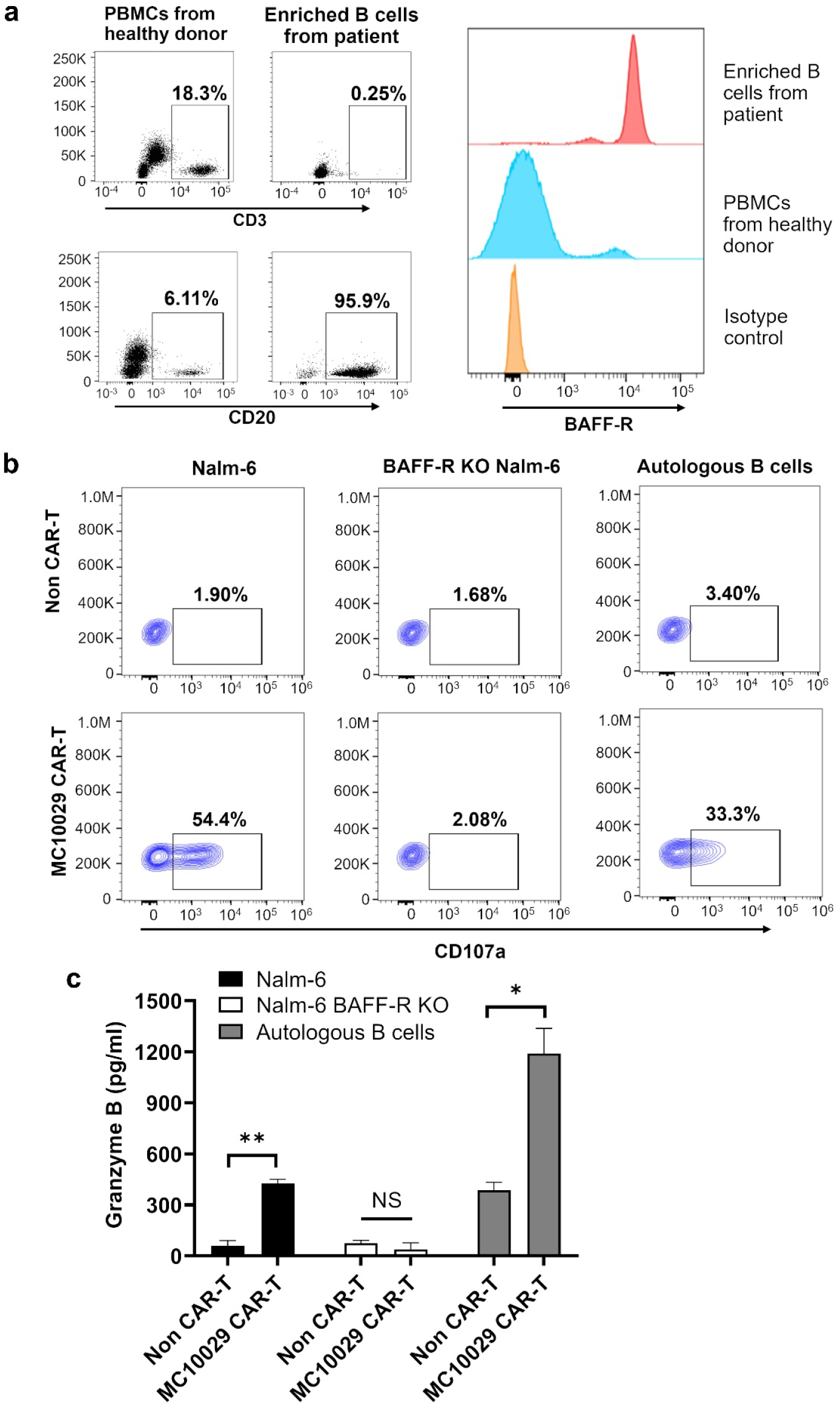
Antigen specific cytotoxicity of patient-derived MC10029 CAR T-cells against autologous B cells. (a) The enrichment process of B cells from the peripheral blood of a representative patient efficiently depletes T cells (CD3 positive cells), resulting in a significant enrichment of CD20-positive B cells (95.9%). These enriched B cells exhibit a robust positive signal for BAFF-R expression (as shown in the top histogram). PBMCs from a healthy donor served as a positive control for T and B cell immunostaining. (b) Antigen-specific cytotoxicity was assessed by measuring the surface expression of CD107a as a marker of degranulation. Isolated autologous B cells from a representative patient activated the patient-derived MC10029 CAR T-cells (MC10029 CAR-T), resulting in degranulation. A pair of BAFF-R-positive Nalm-6 and BAFF-R-deficient Nalm-6 cells (Nalm-6 BAFF-R KO) were included to validate the antigen-specific functionality of the CAR T-cells. (c) Incubation of MC10029 CAR T-cells with autologous B cells expressing BAFF-R resulted in a significant release of the cytolytic protein granzyme B, in comparison to Non CAR T-cells (P =0.0123). (* P < 0.05 ; ** P < 0.01; NS=not significant).

### Evaluation of patient-derived MC10029 CAR T-cells against autologous B cells

We expanded our patient analysis to include five additional patients with diverse demographics and medical histories (**Table 1**). The MC10029 CAR T-cells derived from these patients consistently exhibited degranulation activities against their autologous B cells, as evidenced by the release of cytotoxic granules (granzyme A, granzyme B, and perforin). To account for variations in baseline level of T cell activation among patients, we calculated the fold change between MC10029 CAR T-cells and non-CAR T-cells, thereby normalizing the data for inter-patient comparisons (**Table 1**). Due to limited sample availability, we prioritized the use of patients’ B cells for the granule release experiment over the CD107a degranulation assay. The CD107a degranulation data strongly support the cytotoxicity of CAR T-cells against autologous B cells, as all six patients’ MC10029 CAR T-cells consistently showed positive staining for CD107a, indicating their degranulation activity in response to autologous B cells (**Table 1**). The engineered MC10029 CAR T-cells for each patient were confirmed for their antigen-specific functionality using Nalm-6 and BAFF-R-deficient Nalm-6 cells (**Supplementary Figure S2**). In cases where there were insufficient patient B cells (Patient 2, 5, and 6), we utilized MC10029 CAR T-cells against BAFF-R-deficient Nalm-6 as a control to evaluate the cytotoxicity of CAR T-cells against autologous B cells. All patient-derived MC10029 CAR T-cells generated in this study successfully passed our validated product release testing (**Supplementary Table S1**).

## DISCUSSION

Anti-HLA donor specific antibodies have been extensively documented for their critical role in transplant rejection and resulting adverse outcomes.^8^ Considering the role of B cells in generating anti-HLA antibodies in presensitized patients, we propose that the depletion of B cells using CAR T-cells could effectively achieve a sustained remission in anti-HLA antibody production. Consequently, this approach holds the potential to enhance the likelihood of highly sensitized candidates receiving a transplant with a negative cross match, reducing the risk of rejection, and minimizing their wait-time on the transplant waiting list. To support this hypothesis, we utilized our well-established MC10029 CAR that targets BAFF-R on B cells. Leveraging our unique technical platform, which allows for the generation of patient-derived CAR T-cells from minimal blood samples, we successfully assessed the effectiveness of MC10029 CAR T-cells against autologous B cells. Our data clearly demonstrate the cytotoxicity of MC10029 CAR T-cells against autologous B cells obtained from sensitized patients who have experienced failed kidney transplants. While our current experimental system imposes limitations on evaluating changes in alloantibodies, considering the pivotal role of B cells in antibody production, we anticipate a reduction in HLA-specific alloantibodies following CAR T-cell treatment in our forthcoming clinical study. Our optimism is bolstered by a recent clinical study revealing a remarkable decrease in autoreactive antibodies in lupus patients treated with CD19 CAR T-cell therapy.^9^

CAR T-cell therapy offers a notable advantage over monoclonal antibodies with regards to its enduring therapeutic impact on inhibiting alloantibody production, which is crucial for the successful outcome of organ transplants in sensitized patients. We have recently obtained FDA approval of an Investigational New Drug application for MC10029 CAR T-cell therapy in B-cell hematological diseases. This significant milestone paves the way for the pioneering launch of a human clinical trial, marking the first-ever application of CAR T-cell therapy in sensitized patients waiting for life-saving organ transplants.

## Acknowledgements

This project is supported by the “Mayo Clinic Florida CAR-T Manufacturing Program” to HQ, the “Mayo Clinic President’s Discovery Translation Program” to HQ. and MA K-D, and “Mayo Clinic Transplant Benefactor Award” to TJ and SM.

## Data Statement

Specific data related to the patients who provided blood samples presented in this article cannot be shared publicly because of concerns regarding the privacy of participating individuals. The data generated using patient samples will be shared upon reasonable request to the corresponding authors.

## Author contributions

TJ, YL, MA K-D, and HQ created the study concept and designed the studies; YL, YQ, TH, and RMA performed experiments and conducted data analysis; TJ, RMA, SM, and BCT performed all clinical activities; YL and MEG prepared figures; TJ, YL, MEG, HM, MA K-D, and HQ contributed to manuscript writing and editing.

## Supplementary Material

### Supplementary Methods

**Supplementary Figure S1**. Schematic diagram of generating CAR T-cells from patient blood samples.

**Supplementary Figure S2**. Confirmation of antigen-specific functionality of patient-derived MC10029 CAR T-cells.

**Supplementary Table S1**. Final characteristics of the non-CAR T and MC10029 CAR T-cells generated from T cells isolated from patient blood samples.

**Supplementary References**

