## Supplementary figures for "CAR T-cell therapy - Paving the way for sensitized kidney transplant patients"

### Table of Contents

1. **Supplementary Methods**
   Cell lines
   Human blood samples
   PBMCs isolation from patients' blood samples
   T cells and B cells isolation from patients' PBMCs
   Degranulation assays
   Granule release assay
   Statistical analysis

2. **Supplementary Data**
   Supplementary Figure S1. Schematic diagram of generating CAR T-cells from patient blood samples.
   Supplementary Figure S2. Confirmation of antigen-specific functionality of patient-derived MC10029 CAR T-cells.
   Supplementary Table S1. Final characteristics of the non-CAR T and MC10029 CAR T-cells generated from patients' peripheral blood T cells.

3. **Supplementary References**

### 1. Supplementary methods

#### Cell lines

The cell line of Nalm-6 was purchased from Deutsche Sammlung von Mikroorganismen und Zellkulturen GmbH (DSMZ, Germany), and the BAFF-R knockout variant (BAFF-R KO Nalm-6) was generated as previously described.[S1] 293FT and Jurkat cells were obtained from the ATCC. Prior to cryopreservation, the cell lines underwent authentication for the desired antigen using flow cytometry.

#### Human blood samples

Peripheral blood mononuclear cells (PBMCs) from healthy volunteer donors were isolated via leukapheresis using Leukocyte Reduction System (LRS) cones, by the Division of Transfusion Medicine, Mayo Clinic, Rochester, Minnesota, according to current regulatory requirements and as previously described.[S2]

The patients' blood procurement was performed part of a biorepository protocol approved by the Institutional Review Board (IRB) of Mayo Clinic, Florida (IRB# 22-007826). All patients provided written informed consent, and the protocol adhered to the ethical principles of the Declaration of Helsinki. Approximately 40 ml of blood were collected from consenting patients with two draws. Disease-characteristics of these patients were recorded.

#### PBMCs isolation from patients' blood samples

The collected peripheral blood from subjects is diluted 1:1 v/v with PBS and overlaid the above diluted blood sample (20 ml) on the top of 10ml Ficoll (Sigma). The gradient is centrifuged for 20 minutes with 2000 rpm at room temperature. Subsequently, the PBMC layer was carefully separated and isolated. To assess the number and viability of PBMCs, a cell counter (TC 20 Automated Cell Counter, Bio-Rad) was utilized.

#### T cells and B cells isolation from patients' PBMCs

The PBMCs were subjected to the isolation of T cells using the Pan T cell isolation kit (Miltenyi Biotec, Germany), following the instructions provided. In brief, the cells were initially labeled with Biotin-Antibody cocktail and then with MicroBead Cocktail in FACS buffer. Next, the cell suspension was applied onto the LD column for magnetic cell separation. The unlabeled cells that passed through the column were collected as T cells for CAR T-cell production.

The B cells were isolated from PBMCs using the EasySep™ Direct Human B cell isolation kit (STEMCELL technologies, Canada), according to the provided instructions. In summary, the Isolation Cocktail was added to the sample, followed by the addition of RapidSpheres™ for incubation. Subsequently, the tube was placed into the magnet for negative selection. The isolated B cells were stained with anti-CD3 BV605 (BD Biosciences), anti-CD20 BUV395 (BD Biosciences), and anti-BAFF-R-AF647(BD Biosciences) for characterization.

A second-generation BAFF-R-CAR(MC10029) was generated consisting of a novel BAFF-R antibody scFv, IgG4 transmembrane, CD28, and CD3ζ intracellular signaling domains with truncated EGFR.[S1] The CAR cDNA was cloned into pHIV.7 lentiviral vector. Lentiviruses were produced in 293FT cells, concentrated, and titered with Jurkat cells. T isolated from patients' PBMCs were divided into two aliquots. One aliquot advanced to be expanded as non-transduced (non-CAR) T cells whereas the remaining cells advanced to CAR T-cell production. To generate patient-derived CAR T-cells, T cells were activated with Human T-Activator CD3/CD28 beads (Life Technologies) for 24 hours followed by transduction with CAR lentivirus at multiplicity of infection (MOI) = 1; the CAR T-cells were further activated with CD3/CD28 bead stimulation for six days after which the beads are removed, and the CAR T-cells are permitted to expand for an additional seven days. Non-CAR T-cells are non-transduced T cells from the same donor, expanded following the CAR T-cell protocol, and used as a control. Each batch of CAR T-cells were evaluated for cell quality with Fold Expansion and Viability (as determined by Trypan Blue staining) and CAR T-cell specific characterization with Identity (CD3 positive cells) and Potency (EGFR positive T cells) by flow cytometry.

**Degranulation assays**

CAR T-cells were incubated with target cells at an effector-to-target (E:T) ratio of 2:1 in complete RPMI 1640 medium containing GolgiStop Protein Transport Inhibitor Reagent (BD Bioscience) and CD107a APC antibody (BD Biosciences) for 6 hours.[S1] The cells were subsequently stained with anti-CD3 BV605 (BD Biosciences), anti-CD4 PE-Cy7 (BD Biosciences), anti-CD8 APC-Cy7 (BD Biosciences) and anti-EGFR BV421(BD Biosciences). Samples were run on Attune flow cytometer (Thermo Fisher Scientific) or Fortessa flow cytometer (BD Biosciences) and analyzed using FlowJoTM Version 10 software. Non-CAR T-cells from the same patient were used as negative controls.

**Granule release assay**

CAR T-cells and target cells were co-incubated for 72 hours at an E:T ratio of 4:1. After the incubation period, the supernatant was collected and evaluated for granule release. The levels of granule proteins involved in cytotoxic activity, such as granzyme B, granzyme A and perforin, were quantified using a customized U-PLEX Human ELISA kit from Meso Scale Diagnostics, following the manufacturer's instructions (Rockville, MD, USA). This multiplex kit enables the simultaneous measurement of multiple secretory proteins associated with CAR T-cell function.

**Statistical analysis**

All statistical analyses were performed with the GraphPad Prism software. Data are reported as means ± SD and analyzed by a student's *t* test.

### 2. Supplementary Data

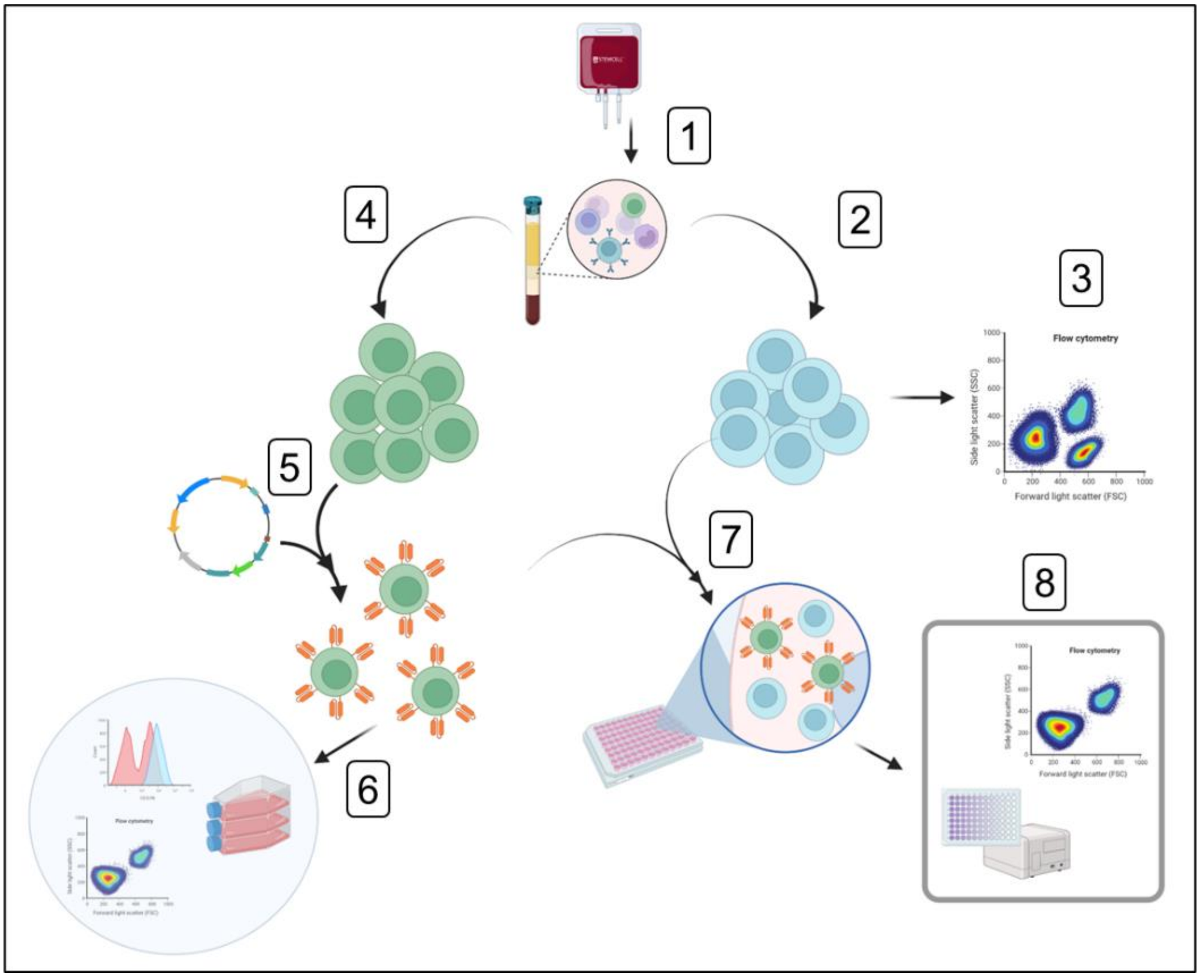

**Supplementary Figure S1. Schematic diagram of generating CAR T-cells from patient blood samples.** (1) Blood is collected from sensitized kidney transplant patient, PBMCs are isolated. (2) B cells are enriched using the EasySep$^{TM}$ Direct Human B cell isolation kit, and surface marker characterization occurs to confirm BAFF-R (3). (4) T cells are isolated using the Pan T cell isolation kit and transduced with MC10029 CAR virus (5). (6) The resulting MC10029 CAR T-cells are expanded, characterized, and subjected to a complement of product release assays. (7) The MC10029 CAR T-cells from the patient are incubated with their autologous B cells. Antigen specific cytotoxicity is determined using flow cytometry (CD107a degranulation assay) and by ELISA (cytokine release assay). Illustration was created with BioRender.com.

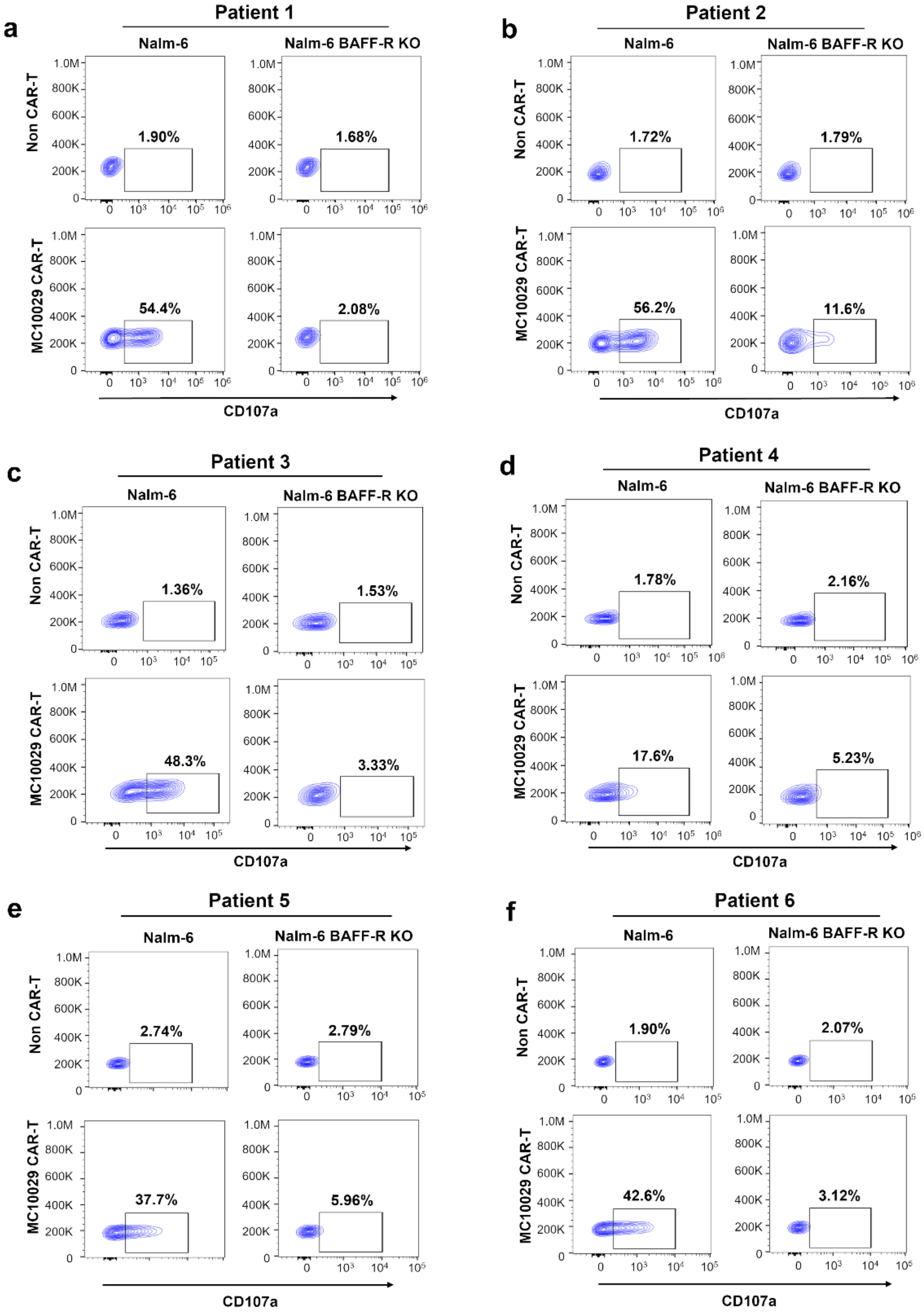
a
Patient 1
Nalm-6
Nalm-6 BAFF-R KO
Non CAR-T
MC10029 CAR-T
1.90%
1.68%
54.4%
2.08%
CD107a
b
Patient 2
Nalm-6
Nalm-6 BAFF-R KO
Non CAR-T
MC10029 CAR-T
1.72%
1.79%
56.2%
11.6%
CD107a
c
Patient 3
Nalm-6
Nalm-6 BAFF-R KO
Non CAR-T
MC10029 CAR-T
1.36%
1.53%
48.3%
3.33%
CD107a
d
Patient 4
Nalm-6
Nalm-6 BAFF-R KO
Non CAR-T
MC10029 CAR-T
1.78%
2.16%
17.6%
5.23%
CD107a
e
Patient 5
Nalm-6
Nalm-6 BAFF-R KO
Non CAR-T
MC10029 CAR-T
2.74%
2.79%
37.7%
5.96%
CD107a
f
Patient 6
Nalm-6
Nalm-6 BAFF-R KO
Non CAR-T
MC10029 CAR-T
1.90%
2.07%
42.6%
3.12%
CD107a

**Supplementary Figure S2. Confirmation of antigen-specific functionality of patient-derived MC10029 CAR T-cells.** Patient-derived MC10029 CAR T-cells were stimulated with BAFF-R positive Nalm-6 cells, resulting in degranulation. Degranulation was measured by evaluating the surface expression of CD107a, a well-established marker of degranulation. BAFF-R-deficient Nalm-6 cells (Nalm-6 BAFF-R KO) were used as an antigen-negative control to confirm the antigen-specific functionality of the CAR T-cells.

**Supplementary Table S1. Final characteristics of the non-CAR T and MC10029 CAR T-cells generated from patients' peripheral blood T cells.**

| | Patient 1 | | Patient 2[a] | | Patient 3 | | Patient 4 | | Patient 5 | | Patient 6 | |
|---|---|---|---|---|---|---|---|---|---|---|---|---|
| | Non CAR-T | CAR-T[b] | Non CAR-T | CAR-T[b] | Non CAR-T | CAR-T[b] | Non CAR-T | CAR-T[b] | Non CAR-T | CAR-T[b] | Non CAR-T | CAR-T[b] |
| **Fold Expansion[c]** | 90.3 | 112 | 81.6 | 74.7 | 58 | 70 | 100 | 98 | 121 | 145 | 90 | 89 |
| **Viability (%) ≥ 70% at D14[d]** | 85 | 83 | 94.3 | 95 | 92.9 | 94.0 | 97.0 | 96.0 | 94.5 | 90.7 | 92.8 | 93.7 |
| **Identity (%) ≥ 80%[e]** | 99.7 | 99.5 | 96.6 | 97.5 | 99.7 | 99.7 | 97.8 | 98.2 | 99.1 | 99.2 | 99.3 | 99.4 |
| **Potency (%) ≥ 10%[f]** | 0.71 | 28.1 | 0.57 | 25.1 | 0.91 | 28.2 | 0.46 | 27.9 | 0.79 | 28.5 | 0.83 | 20.3 |

**T cells isolated from patient blood samples.**

[a] Patient 2 received prior treatment with rituximab in 2020 and cyclophosphamide until 2022.

[b] CAR-T = MC10029 CAR T-cells

[c] Fold expansion is a parameter that quantifies the increase in cell numbers during the 14-day CAR T-cell process, starting from the original 1 million cells.

[d] Viability is measured on day 14 (D14) with the acceptance criteria of ≥ 70%.

[e] Identity is measured on day 14 using a CD3 antibody with the acceptance criteria of ≥ 80% cells having the characteristic CD3 surface marker.

[f] Potency is measured on day 14 using an EGFR antibody with the acceptance criteria of ≥ 10% cells having the tEGFR surface marker, a proxy marker for the CAR T-cells. tEGFR is an inactive surface protein that is expressed as part of the same vector as MC10029 CAR.
